# Functional heterogeneity of beta bursts in childhood reveals a dimensional neural signature of motor skill

**DOI:** 10.1101/2025.09.29.679222

**Authors:** Holly Rayson, Sarah A. Gerson, Jennifer Keating, Catherine R.G. Jones, Catherine Purcell, James J Bonaiuto, Ross E Vanderwert

**Affiliations:** Marc Jeannerod Institute of Cognitive Sciences, ISC, CNRS UMR5229, Lyon, France; Université Claude Bernard Lyon 1, Université de Lyon, France; Hospices Civils de Lyon, Lyon, France; CHU de Clermont-Ferrand, Clermont-Ferrand, France & CH Montluçon-Néris les Bains; School of Psychology, Cardiff University Centre for Human Developmental Science (CUCHDS), Cardiff, United Kingdom; School of Social Sciences, Cardiff University, Cardiff, United Kingdom; School of Healthcare Sciences, Cardiff University, Cardiff, UK; School of Psychology, Cardiff University Brain Research Imaging Centre (CUBRIC), Cardiff, United Kingdom

**Keywords:** beta bursts, motor control, developmental coordination disorder, EEG, sensorimotor cortex, motor development

## Abstract

Activity in the beta frequency band appears to play a critical role in sensorimotor processing, with transient ‘bursts’ exhibiting distinct waveform motifs that may reflect different neural computations. However, how these dynamics relate to motor development and neurodevelopmental conditions remains unclear. Here, we used EEG to investigate sensorimotor beta bursts in children with and without developmental coordination disorder (DCD). We characterized beta bursts during action execution and observation of gross and fine motor actions. Conventional spectral features and overall burst rates did not differ between groups. In contrast, children with DCD showed altered rate modulation of specific burst waveform motifs, including atypical hemispheric lateralization and disrupted interhemispheric connectivity. Critically, motif-specific burst rates were systematically related to motor performance across individuals, with similar relations observed across groups. These findings indicate that waveform-specific beta burst dynamics track graded variation in motor performance, rather than mapping selectively onto diagnostic status. Beta burst variability may therefore reflect the functional expression of motor performance, rather than a disorder-specific neural mechanism underlying DCD. This work highlights the importance of beta burst waveform diversity for understanding variability in sensorimotor function across development.

## 1. Introduction

Sensorimotor beta activity (13-30 Hz in adults) has long been associated with motor control and is traditionally conceptualized as a sustained oscillatory rhythm that modulates in power prior to, during, and after movement (Pfurtscheller, 1981; Pfurtscheller et al., 1996). However, more recent work indicates that beta activity occurs in transient, burst-like events, rather than as a continuous oscillation (Feingold et al., 2015; Little et al., 2019; Sherman et al., 2016). These *beta bursts* are temporally sparse and behaviorally relevant, predicting response time, motor errors, and stopping performance in adults (Little et al., 2019; Wessel, 2020; West et al., 2023), and are modulated not only during action execution, but also during observation of others’ movements (Rayson et al., 2023), implicating them in motor simulation and internal model engagement (Jeannerod, 2001; Press et al., 2011). Developmental work further suggests that beta activity occurs as bursts from infancy, with age-related shifts in peak frequency, timing, duration, and hemispheric lateralization (Cheyne et al., 2014; Gaetz et al., 2010; Rayson et al., 2022, 2023; Rier et al., 2024; Wilkinson et al., 2024). These findings indicate that maturation of sensorimotor function may involve progressive refinement of transient beta events rather than simple changes in sustained oscillatory power. Although the precise computational role of beta bursts remains unresolved, their developmental tuning and behavioral relevance suggest that burst dynamics provide a useful framework for understanding how transient beta activity contributes to sensorimotor processing (Rayson et al., 2025).

Most studies implicitly treat beta bursts as homogenous events, yet recent work demonstrates that individual bursts vary systematically in their time-domain waveform shape. Rather than defining burst types using spectral features (e.g., peak frequency, power, or duration), waveform-based approaches represent each burst as a vector of voltage (or magnetic field strength in the case of magnetoencephalography) across time and apply dimensionality reduction (e.g., PCA) to identify axes of shape variability (Agouram et al., 2025; Papadopoulos et al., 2024; Rayson et al., 2023; Szul et al., 2023). A “motif” in this context refers to the characteristic waveform shape at the extreme ends of a given principal component, that is, bursts with high positive or negative loadings that share reproducible configurations of peaks, troughs, asymmetries, or temporal skew relative to the mean burst waveform. Motifs are therefore not discrete physiological generators, but statistical patterns of shape variation within the burst population. These motifs differ in their rate dynamics and lateralization, and accumulating evidence suggests that they index distinct functional processes (Agouram et al., 2025; Rayson et al., 2023; Szul et al., 2023) such as behavioural error (Moreau et al., 2026).

Developmental findings support the functional relevance of this variability. Infant bursts share a similar canonical waveform with adults but are temporally longer; after accounting for duration differences, their waveforms align closely in shape (Rayson et al., 2023). Moreover, variability around the mean waveform is structured and conserved across age, while motif-specific rate modulation becomes increasingly precise from infancy to adulthood (Rayson et al., 2023), consistent with progressive specialization of sensorimotor computations. Examining waveform-defined motifs across children spanning a range of motor abilities, including those with developmental coordination disorder (Wilson et al., 2017) therefore provides a principled framework for testing whether specific dimensions of burst shape variability track individual differences in motor skill and interhemispheric organization.

Recent evidence indicates that beta bursts with distinct waveform shapes exhibit dissociable patterns of rate modulation and hemispheric lateralization during movement, with some motifs showing increasingly precise lateralization across early development (Rayson et al., 2023). These findings suggest that waveform-defined burst types may reflect distinct computational roles within distributed sensorimotor networks, potentially including interhemispheric coordination. Motor ability varies substantially across development, and individual differences in coordination are associated with variability in hemispheric specialization and functional connectivity within sensorimotor systems (Brown-Lum & Zwicker, 2015; McLeod et al., 2016; Querne et al., 2008; Tallet et al., 2013). Studying children with a wide range of motor proficiency, including those meeting criteria for developmental coordination disorder (Wilson et al., 2017), therefore provides a principled way to test whether specific waveform-defined beta burst motifs scale with motor performance and interhemispheric dynamics. While DCD is defined categorically at the clinical level, variation in motor performance and its neural substrates may nonetheless be continuous across individuals. Under this framework, DCD therefore serves as a model of pronounced motor difficulty within a broader continuum, enabling us to examine how variation in burst motif dynamics relates to motor coordination.

Here, we characterized sensorimotor beta burst dynamics in children spanning a broad range of motor abilities during action execution and observation, including children with and without DCD. We explicitly tested for differences between groups in the modulation, lateralization, and interhemispheric connectivity of waveform-defined burst motifs, and additionally examined whether motif-specific burst rates were related to individual differences in motor performance. We show that specific burst motifs display altered rate modulation and hemispheric asymmetry during movement in children with DCD, and that motif-specific burst rates are systematically related to motor performance across individuals. Notably, although these effects differentiate groups, the relationship between burst dynamics and motor performance is continuous across the full sample. These findings suggest that variability in waveform-specific beta burst dynamics reflects graded differences in the organization of interhemispheric sensorimotor communication, along a continuum of motor performance.

## 2. Methods

### 2.1 Participants

Twenty-three 8- to 12-year-old children with DCD (*M* = 10.88, *SD* = 1.56; 18 male) were recruited via social media and the Dyspraxia Foundation, and 33 typically developing children (*M* = 9.85, *SD* = 1.30; 17 male) were recruited via social media as a comparison group. Participants in the DCD group were required to have received a clinical diagnosis of DCD, confirmed for this study via parent report and converted total score on the Movement Assessment Battery for Children 2^nd^ Edition (MABC-2; Henderson et al., 2007) at or below the 16^th^ percentile at the time of the testing session. In line with the Leeds Consensus Statement (2006), participants with a diagnosis of DCD who had co-occurring neurodevelopmental conditions were not excluded from participating, but additional diagnoses were noted. In the final sample, co-occurring conditions were autism (*n* = 3), ADHD (*n* = 1), dyslexia (*n* = 1), and autism, dyslexia, developmental language disorder and learning difficulty (*n* = 1). Children were not recruited to the comparison group if they had a diagnosis of a neurodevelopmental condition or developmental delay.

Nine children from the comparison group were excluded from the analyses characterizing beta burst group differences as they scored below the 25^th^ percentile on the MABC-2. This was to ensure we were not including children in the comparison group who did not have normative motor development (French et al., 2018). A further six participants were excluded during EEG processing due to insufficient data (DCD *n* = 4; comparison *n* = 2). The final sample therefore included 19 children with DCD (*M*_age_ = 11.17 years, *SD* = 1.28; 14 male; 14 right-handed) and 22 children without DCD (*M*_age_ = 9.78 years, *SD* = 1.40; 12 male; 21 right-handed). Full testing and demographic details are available in Keating et al., (2023); however we note here that the comparison group had significantly higher IQ scores (*p* = .011) (assessed by the Weschler Abbreviated Scales of Intelligence, 2nd edition [WASI-II]; Weschler, 2011) and were significantly younger than the DCD group (*p* = .002). Written informed consent was obtained from the participant’s caregiver before the start of the experiment. Each child also provided written assent to take part. Ethical approval was granted by the Ethics Committee at Cardiff University (EC.20.03.10.5977R2). Participants received travel expenses and a small prize.

### 2.2 Movement Assessment Battery for Children

The Movement Assessment Battery for Children 2^nd^ Edition (MABC-2; Henderson et al., 2007) test component was used to assess motor skills. The MABC-2 has good internal consistency and test re-test reliability (Henderson et al., 2007; Wuang et al., 2012). The assessment consists of eight tasks organized into three subsets: manual dexterity, aiming and catching, and static and dynamic balance. Age-adjusted standard scores were calculated for each subtest, and a total difficulty score was created from which percentiles can be derived. A total score below the 5^th^ percentile indicates significant motor difficulties, a score between the 5^th^ and 16^th^ percentile indicates a risk of motor difficulties, and a score above the 25^th^ percentile indicates age-expected motor performance.

### 2.3 Task and Stimuli

Video stimuli for the EEG experiment were filmed using a handheld camera fixed to a tripod. The camera was positioned overlooking the model’s shoulder while she performed the actions to best match the participant’s perspective of the actions as if they were performing them, while also providing a clear view of the entire action. We chose an ego-centric perspective because previous studies have suggested this elicits greater sensorimotor activity (Drew et al., 2015; Fu & Franz, 2014; Jackson et al., 2006).

All videos were trimmed to fit within a 10-second period, with three types of videos in total (gross motor, fine motor, and non-biological movement). The gross motor video displayed a model sitting with the hand starting at rest on the edge of a table, extending the arm to reach out and tap a red square, and then returning the arm to rest. This action took approximately 2 seconds and was repeated 5 times. The fine motor video displayed a hand holding a red pen and tracing a flower figure (based on the Movement Assessment Battery for Children; Henderson et al., 1992). The tracing lasted approximately 20 seconds and was split into two separate videos; therefore, half of the fine motor action observation trials began with an empty flower and showed the hand tracing halfway around and the other half began with the hand already halfway around the flower and continued tracing to the end of the flower. The non-biological movement videos were of kaleidoscope patterns taken from Pixabay (www.pixabay.com). All videos of actions were performed by a right-handed person and were mirrored to make the actions appear to be performed by a left-handed model for left-handed participants (n = 6). All videos were presented on the center of the screen, subtending a visual angle of ∼28° (horizontal) and ∼14° (vertical). Example stimuli are displayed in Figure 1 and are available at https://osf.io/vnup9/.

**Figure 1.**
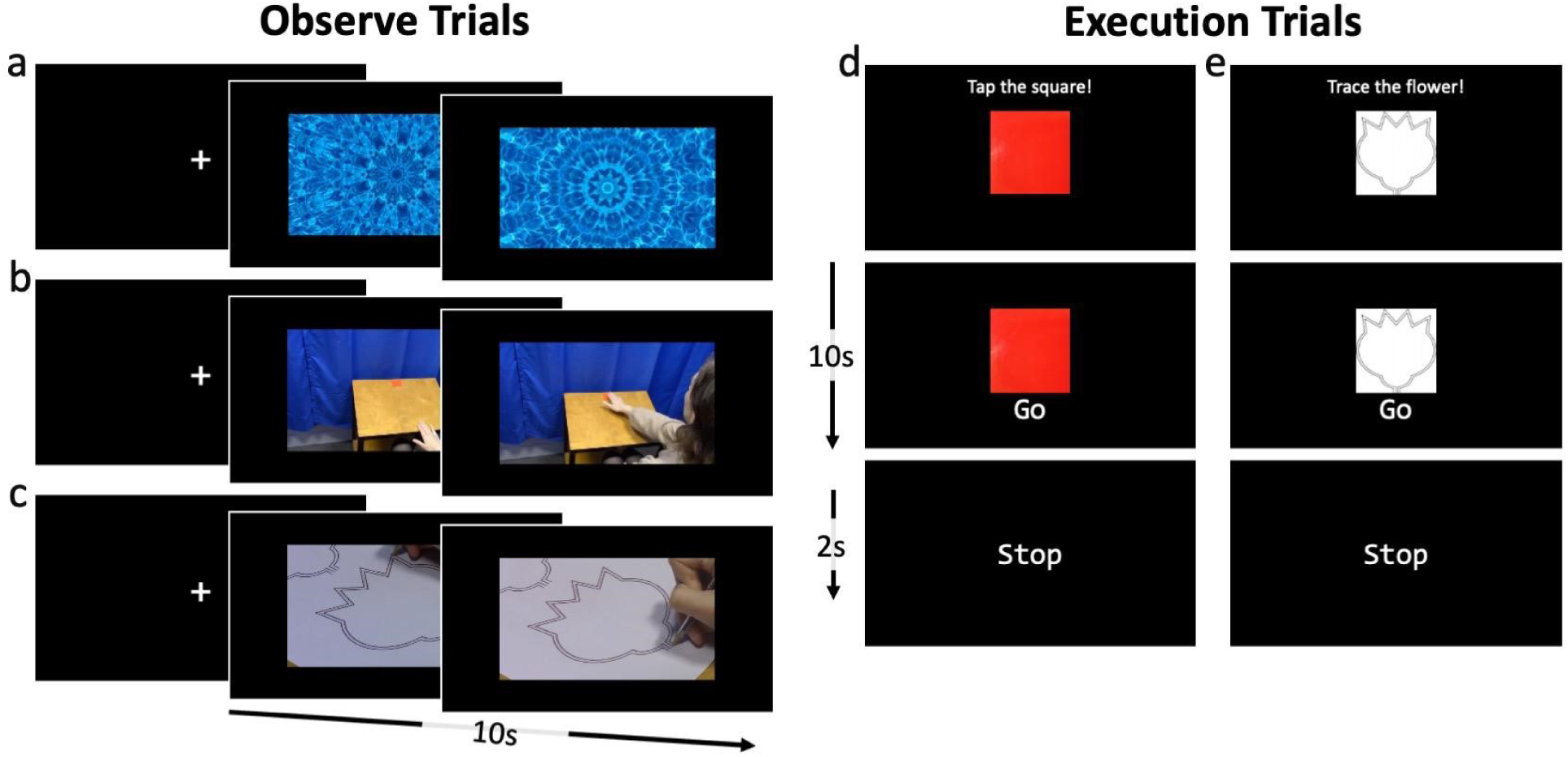
Example Trials in the EEG Task. Observe trials began with a fixation cross to cue participants to remain still and observe the videos for 10 seconds. Example screenshots of the videos are shown for Kaleidoscope (a); the hand reaching for the red square for Gross Motor (b); and a hand tracing a flower for Fine Motor (c) trials. Execution trials began with action instructions to prepare participants; specifically to “Tap the square!” for Gross Motor (d) or “Trace the flower!” for Fine Motor (e) trials. The “Go” screen was then displayed for 10 seconds to initiate participants execution and then the “Stop” screen was displayed for 2 seconds to end the trial.

The EEG task consisted of five conditions adapted from Hobson and Bishop (2016; Figure 1). Two conditions were execution trials: Execution Gross motor (ExG) and Execution Fine motor (ExF). These began with an instruction screen describing the action with an image of the action stimulus (i.e. a red square for ExG/the flower for ExF). Once the participant was ready, the experimenter initiated the trial with a button press. During the trial, the command ‘GO’ was displayed beneath the action stimulus for 10 seconds while participants performed the action. A ‘STOP’ command was then displayed for 2 seconds. Three conditions consisted of observation trials using the videos described above. These began with a fixation cross. Once the participant was ready, the experimenter initiated the trial with a button press. During the trial, either the Observation Gross motor (ObG), Observation Fine motor (ObF), or the non-biological movement (kaleidoscope; Kal) video played for 10 seconds. Participants completed 40 trials of each condition split across two blocks. Trials were presented in a random order and timing was controlled by E-Prime 3.0 software (Psychology Software Tools).

### 2.4 Procedure

Each child was prepared for the EEG acquisition, including placement of the EEG cap and preparation of electrodes with electrogel, and then brought into the testing room. Their caregiver was invited to observe the testing session from an adjoining room while they completed questionnaires on a tablet computer (see Keating et al., 2023 for details). During placement of the EEG cap, the child completed the Edinburgh Handedness Inventory - Short Form (Veale, 2014) to determine whether they were dominantly left- or right-handed. Once the EEG was fully set-up, the task was explained to the participant.

Participants sat at a table with a red square affixed to the surface across from a 17” computer screen approximately 80 cm away. The session began by measuring resting-state EEG as the child sat with their eyes first open for 30 seconds, and then their eyes closed for 30 seconds. This procedure was repeated three times.

Following resting measures, children practiced the tasks via completion of two observation trials (one ObG and one ObF) and two execution trials (one ExG and one ExF). Experimenters ensured children were on-task and did not emphasize speed or accuracy during execution trials. Participants then completed the EEG acquisition, with the entire session lasting around 25 minutes. Once the EEG session was complete, the EEG cap was removed, and the participants completed the MABC-2 and WASI-II (in this fixed order). Children could take breaks during testing as needed, with the whole session lasting approximately 100 minutes.

### 2.5 EEG Acquisition

Continuous EEG was recorded with the BrainVision actiCHamp Plus system (Brain Products GmbH, Gilching, Germany) from 32 channels placed according to the International 10-20 System, and referenced online to the vertex electrode (Cz). ECI Electro-Gel (GVB geliMED GmbH, Germany) was used to reduce impedances, which were kept below 20 kΩ. Data were sampled at 500 Hz. Video recordings of all EEG sessions were coded to identify trials in which participants were not watching a video or completing the required action. Any trials where participants were visibly not attending or were making off-task movements were removed prior to analysis (Observation: *Mdn* = 4, range 0-13 trials; Execution: *Mdn* = 2, range 0-8 trials).

### 2.6 Data preprocessing

EEG data were preprocessed in MATLAB (R2018a) using a custom version of the MADE pipeline (Debnath et al., 2020), augmented with artifact-detection routines from the NEAR pipeline (Kumaravel et al., 2022). Data were high-pass filtered at 1 Hz and low-pass filtered at 100 Hz using EEGLAB (v2021.1; Delorme & Makeig, 2004) Hamming-window FIR filters (high-pass cutoff at 0.5 Hz; low-pass cutoff at 105 Hz, reflecting the transition-band specification in the filter design). Prior to cleaning, discontinuous segments outside the task were removed by trimming the continuous recording to 1.5 s before the first event and 1.5 s after the last event.

Bad channels were identified using NEAR’s local outlier factor (LOF) procedure, with an initial LOF cutoff of 2.5 and adaptive thresholding enabled (the cutoff was incremented when >10% of channels were flagged); flat-line detection was also enabled (5 s tolerance), while the periodogram-based detector was disabled (comparison: 0 - 3 channels, *M* = 0.24, *SD* = 0.66; DCD: 0 - 12 channels, *M* = 0.70, *SD* = 2.51). Artifact Subspace Reconstruction (ASR) was then applied to correct non-stereotyped artifacts (correction mode, burst criterion = 13). Stereotyped artifacts were subsequently removed using ICA, computed on a copy of the dataset segmented into 1 s epochs; epochs exceeding ±1000 µV were rejected prior to ICA, and channels with >20% rejected 1 s epochs were removed from the ICA copy and correspondingly excluded from the original data before transferring ICA weights. Artifactual independent components were identified using the Adjusted-ADJUST plugin with default parameters (Leach et al., 2020; Mognon et al., 2011) and removed from the original continuous dataset (comparison: 1 - 19 components, *M* = 8.03, *SD* = 4.66; DCD: 1 - 14 components, *M* = 7.74, *SD* = 2.83).

Following ICA-based correction, data were epoched into task segments time-locked to condition markers (Kal, ObF, ObG, ExF, ExG) using a -8 to 15 s window. Artifact rejection was performed at the epoch level using a ±150 µV threshold; epochs were rejected if ≥10% of channels exceeded threshold (comparison: 1 - 99 epochs, *M* = 34.24, *SD* = 29.12; DCD: 9 - 99 epochs, *M* = 31.96, *SD* = 27.79), otherwise the affected channels were interpolated within that epoch. After artifact rejection, remaining missing channels were interpolated (spherical interpolation) and data were average re-referenced. Finally, 50 Hz line noise was removed using an iterative Zapline procedure (de Cheveigné, 2020) implemented in the MEEGKit package (https://nbara.github.io/python-meegkit/), using a window size of 10 Hz for polynomial fitting and 2.5 Hz for noise peak removal and interpolation. Participants with fewer than 5 trials for a condition were excluded from analysis of that condition. All source code for preprocessing is available at https://github.com/danclab/dcd_bursts/tree/main/preprocessing.

### 2.7 Data-Driven Definition of the Beta Band

All analyses focused on the C3 and C4 electrodes, which were selected *a priori* because they overlie the left and right sensorimotor cortex, which are the canonical loci of movement-related beta modulation (Pfurtscheller, 1981; Pfurtscheller et al., 1996). To define the frequency limits of the beta band in a data-driven manner, power spectral densities (PSDs) were computed for each participant from 0.5 to 40 Hz using Welch’s method (1 s Hann windows, 50% overlap, 10× zero-padding). PSDs were parameterized using specparam (Donoghue et al., 2020) to separate periodic (oscillatory) and aperiodic components.

Periodic components were averaged across C3 and C4 electrodes and then across participants from both groups. Spectral peaks were identified using an iterative peak-fitting procedure in which the frequency-domain periodic spectrum was fit with a Gaussian at its global maximum, subtracted, and re-evaluated until no additional peaks remained. Peak frequency bands were defined using the full width at half maximum (FWHM) of each Gaussian, retaining only peaks with a minimum FWHM of 1 Hz below 10 Hz or 3 Hz above 10 Hz. This procedure yielded a beta range of 14-31.25 Hz, which was used in all subsequent analyses.

### 2.8 Conventional Beta Power Analysis

To characterize canonical beta modulation, time-frequency (TF) representations of each trial were computed using an adaptive superlet transform (Moca et al., 2021), with varying central frequencies (1-50 Hz) and a fixed number of cycles (4 cycles) within a Gaussian envelope. The wavelet order (i.e., the multiplier for the number of cycles in the wavelet) varied linearly from 1 to 40 over the frequency range. Time-frequency representations were averaged across the defined beta band and baseline-corrected relative to -7 to -1 s using percent signal change. Group-level time courses were compared using cluster-based permutation testing (Maris & Oostenveld, 2007), with between-group comparisons performed via independent-samples cluster tests and within-group modulation assessed using one-sample cluster tests against zero.

To complement time-resolved analyses, beta power was averaged within predefined temporal windows (baseline: -7 to -1 s; task: 1-9 s; post-task: 11-14 s), and analyzed using linear mixed-effects models to assess differences across groups and task windows within each hemisphere and execution condition. For each hemisphere (contralateral, ipsilateral) and execution condition (ExF, ExG), window-averaged beta power was modeled as a function of group (DCD, comparison), window, and their interaction, with participant included as a random intercept. Models were fit separately for each hemisphere and condition. Statistical significance of fixed effects was evaluated using Type III Wald χ² tests (car v3.1.0; Fox & Weisberg, 2018). Where appropriate, pairwise post hoc comparisons were performed using Tukey-corrected estimated marginal means (lsmeans v2.30; Lenth et al., 2020). All models were fit in R (v4.2.2; R Core Team, 2022) using the lme4 package (v1.1.30; Bates et al., 2014).

### 2.9 Lagged Hilbert Autocoherence

To assess whether beta activity reflected sustained rhythmic oscillations or transient burst-like dynamics, we computed lagged Hilbert autocoherence (LHaC) over the 1-40 Hz range during the 0-10 s execution interval. LHaC quantifies the degree to which oscillatory amplitude fluctuations remain temporally coherent across lags; lower values in the beta band relative to neighboring frequencies indicate reduced sustained rhythmicity and greater burst-like structure (Zhang et al., 2025). For each participant and hemisphere, LHaC was computed across temporal lags (0.1-4.5 cycles). Group-level LHaC spectra were obtained by averaging across participants within hemisphere.

### 2.10 Beta Burst Detection Algorithm

The lagged Hilbert autocoherence analysis indicated that beta activity over sensorimotor cortex was weakly rhythmic and therefore better characterized as transient burst events rather than sustained oscillations. We therefore analyzed beta activity at the level of individual burst events using a recently developed adaptive burst detection algorithm (Rayson et al., 2023; Szul et al., 2023) designed to capture transient oscillatory events over a wide amplitude range without relying on a fixed global threshold (Figure 2). For each trial, time-frequency (TF) representations were computed using the superlet transform (Moca et al., 2021), which optimally balances time and frequency resolution, and is thus particularly suited for detecting transient bursts within narrow frequency bands (Figure 2a). As for conventional power analyses, an adaptive superlet transform was used, with varying central frequencies (1-50 Hz) and a fixed number of cycles (4 cycles) within a Gaussian envelope. The wavelet order (i.e., the multiplier for the number of cycles in the wavelet) varied linearly from 1 to 40 over the frequency range.

**Figure 2.**
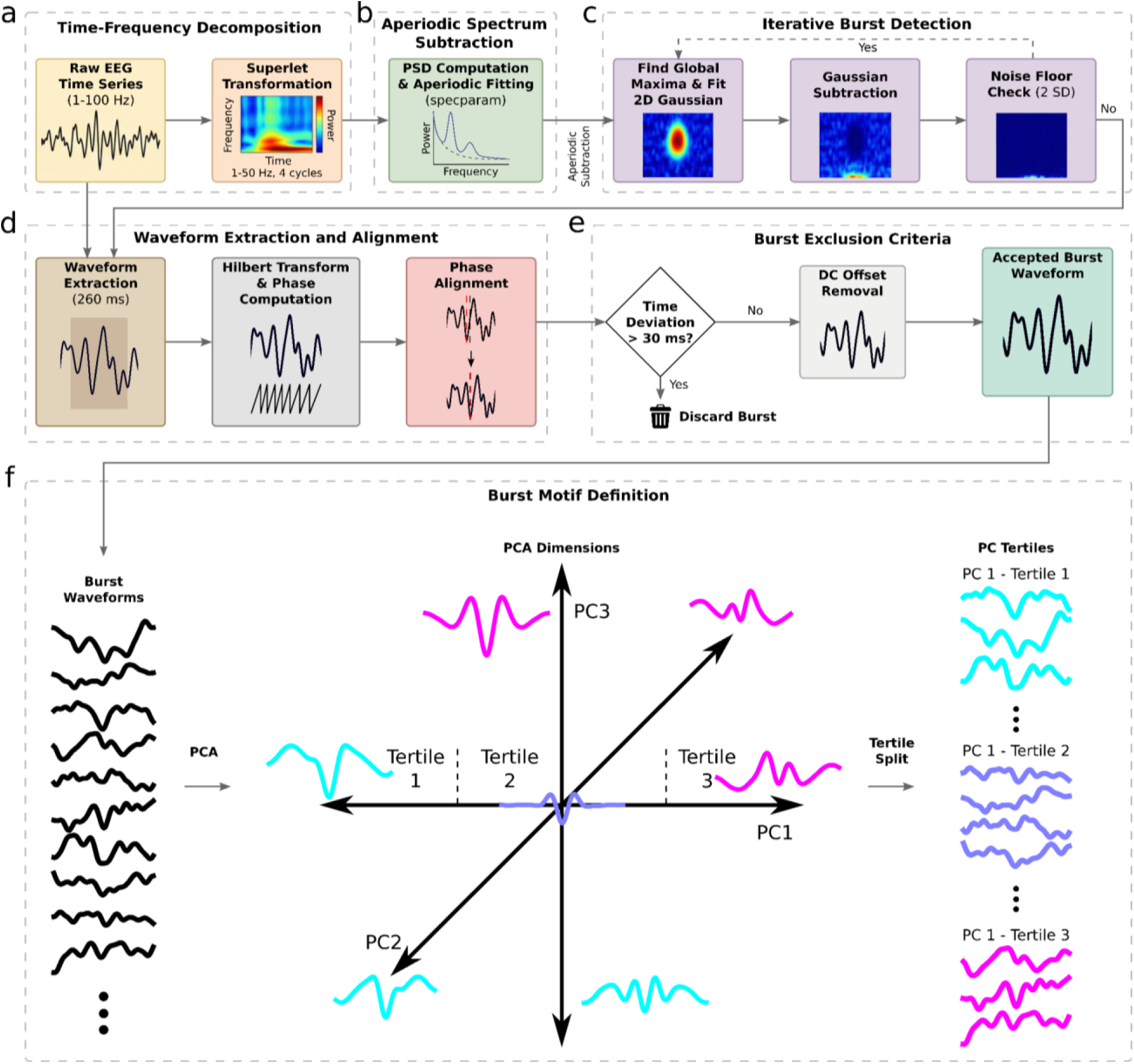
Overview of the beta burst detection, waveform extraction, and waveform-based analysis pipeline. (a) Raw EEG time series are transformed into the time-frequency domain using the superlet transform. (b) Power spectral densities are computed and parameterized to separate periodic and aperiodic components, allowing subtraction of the aperiodic fit from the TF representation. (c) Burst detection proceeds iteratively by identifying the global TF maximum, fitting a two-dimensional Gaussian, subtracting it from the residual TF matrix, and repeating until no peaks exceed a noise threshold (2 SD). (d) For each detected burst, a 260 ms time-domain waveform is extracted, phase-aligned using the Hilbert transform, (e) checked for temporal consistency with the TF peak, DC-corrected, and retained for further analysis. (f) Bursts are projected onto selected principal components (e.g., PC6 and PC8), stratified into tertiles according to their PC scores, and grouped into waveform motifs for subsequent comparisons of burst dynamics across experimental conditions and participant groups.

To isolate oscillatory burst activity, PSDs were derived from each trial’s TF decomposition and parameterized using specparam to estimate the aperiodic (1/f-like) component of the spectrum (Figure 2b). This step is critical because aperiodic activity contributes broadband power that can vary across frequency and time, and if left unaccounted for, can bias burst detection by inflating apparent oscillatory peaks or distorting their frequency and duration estimates. The aperiodic fit was therefore subtracted from the TF representation, yielding a residual TF matrix that more selectivity reflects band-limited power (Brady & Bardouille, 2022). Burst detection was then performed directly in this residual TF space (Figure 2c). At each iteration, the algorithm identified the global maximum in the residual TF matrix, defined jointly over the time and frequency dimensions. A two-dimensional Gaussian was then fit to this peak, with one dimension capturing the burst’s temporal extent (duration) and the other capturing its spectral extent (frequency span).

The fitted Gaussian provides estimates of peak time, duration, peak frequency, frequency span, and peak amplitude, and was subsequently subtracted from the residual TF matrix. The algorithm then proceeded to the next iteration on the updated residual. Iteration continued until no remaining TF peak exceeded a noise floor defined as 2 standard deviations above the mean residual power across all time and frequency bins, recomputed in every iteration. To minimize edge effects, burst detection was applied to TF data spanning 11-34.25 Hz, and only bursts with peak frequencies within the predefined beta band (14-31.25 Hz) were retained for further analysis.

For each detected burst, the corresponding time-domain waveform was extracted from the minimally filtered EEG signal (1 Hz high-pass, 100 Hz low-pass) using a 260 ms window centered on the TF peak time (Figure 2d; Szul et al., 2023). To ensure consistent alignment of waveform features across bursts, waveforms were phase-aligned by band-pass filtering each burst within its detected frequency span, computing the instantaneous phase using the Hilbert transform, and re-centering the unfiltered waveform on the phase minimum closest to the TF peak (Boto et al., 2022). Bursts were excluded if the phase-aligned peak deviated by more than 30 ms from the TF-detected peak time (Figure 2e). Finally, the DC offset was removed to yield the accepted burst waveform used in subsequent analyses. All code for the burst detection algorithm is available at https://github.com/danclab/burst_detection.

### 2.8 Burst analysis

For each detected burst, the iterative burst detection algorithm described above estimated peak amplitude, peak frequency, duration, and frequency span from the fitted two-dimensional Gaussian in the residual time-frequency representation. We first compared beta burst features (duration, peak amplitude, peak frequency, and frequency span) between the comparison and DCD groups to ensure that any subsequent group differences were not driven by conventional time-frequency characteristics. Statistical significance was assessed using nonparametric permutation tests (N = 100,000), in which group labels were randomly shuffled and the observed difference in mean feature values was evaluated against the resulting null distribution.

Overall burst rate was then computed separately for contralateral and ipsilateral sensorimotor electrodes during fine (ExF) and gross (ExG) motor execution. For each participant and condition, burst occurrences were binned in 0.2 s intervals from -8 to 15 s, converted to rates (Hz), and normalized by the number of trials. The resulting time series were smoothed with a Gaussian kernel (σ = 2 bins) and baseline-corrected by subtracting the mean rate during the pre-task interval (-7 to -1 s). Group differences in burst-rate dynamics were evaluated using cluster-based permutation testing (1,000 permutations) on the baseline-corrected time series, controlling for multiple comparisons across time.

We then used Principal Component Analysis (PCA) to characterize variability in burst waveform shape and to determine whether this variability differed between the comparison and DCD groups (Figure 2f). Unlike conventional time-frequency features (e.g., peak frequency or amplitude), which are derived from spectral representations, waveform analysis operates directly on the raw time-domain signal of each detected burst. PCA identifies orthogonal axes that capture the dominant dimensions of waveform variability across bursts. These components represent statistical directions of variance and are not assumed to correspond to discrete physiological generators. Throughout the manuscript, we use the term “motif” as a descriptive shorthand for the waveform shapes associated with extreme loadings on a given component.

Burst waveforms were extracted during the detection procedure and pooled across conditions for PCA. We specified 20 components to ensure that the decomposition captured the full range of observable waveform variability. This dimensionality was chosen conservatively: across multiple datasets analyzed with this framework, permutation-based significance testing has consistently identified no more than approximately 10-12 components as exceeding chance levels of explained variance (Agouram et al., 2025; Papadopoulos et al., 2024; Rayson et al., 2023; Szul et al., 2023). Specifying 20 components therefore provides a stable upper bound that exceeds the number of meaningful components without constraining the solution a priori.

We first conducted separate PCAs for each group (after robust scaling), treating each time point as a feature and defining the mean burst waveform as the origin of a 20-dimensional space. Each burst was represented by its scores along each principal component, indicating its deviation from the mean waveform along that dimension (Figure 2f). To assess whether the structure of waveform variability differed between groups, we compared eigenvectors using Spearman correlation and observed high correspondence across groups. A single global PCA was therefore fit to all bursts pooled across participants, enabling direct comparison within a shared representational space.

To determine which components captured meaningful waveform variance, we compared the variance explained by each component in the real data with null distributions obtained from 100 iterations of temporally shuffled data, in which time points within each waveform were permuted to destroy temporal structure while preserving marginal distributions (Vieira, 2012). P-values were defined as the proportion of shuffled solutions explaining greater variance than the empirical component. After Bonferroni correction (α = 0.0025), PCs 1-11 were retained for further analysis.

To identify waveform motifs that were modulated during movement, we compared principal component (PC) scores between baseline (-7 to -1 s) and task (1 to 9 s) windows within each execution condition (ExF, ExG). For each subject, bursts were assigned to 100 ms time bins, and mean PC scores were computed per bin, normalized by trial count, and smoothed (Gaussian kernel, σ = 2.5 bins). This yielded a time-resolved estimate of the average contribution of each PC to the burst population over the course of the trial. For each PC, we then computed the difference between baseline and task window means separately for each subject and group. Statistical significance was assessed using paired permutation tests (sign-flipping across subjects; 1,000 permutations). PCs showing a significant baseline-task difference (*p* < 0.05) were considered task-modulated. This analysis identified PCs 3, 6, and 8 as significantly modulated during execution. Because changes in mean PC score may reflect altered prevalence of bursts at different positions along that dimension rather than purely within-burst shape changes, we next quantified waveform-specific burst rates by stratifying bursts according to their PC score and examining condition-dependent changes in their occurrence.

Importantly, because principal components define continuous dimensions of waveform variation, group differences are not expected to manifest uniformly across all portions of a component’s score distribution; instead, they may appear selectively at one extreme of that dimension, reflecting a redistribution in the prevalence of bursts with particularly high or low loadings rather than a global shift in mean waveform shape. Therefore, for each PC that exhibited task-related modulation, we quantified waveform-specific burst rate by stratifying bursts into tertiles of PC score (based on the global percentiles of that PC across all bursts; Figure 2f). Burst rates were then estimated separately for contralateral and ipsilateral sensorimotor electrodes and for each execution condition (ExF, ExG). Within each subject, bursts were binned in 200 ms intervals over the full trial epoch (-8 to 15 s), converted to rates (Hz) by dividing by bin width, and smoothed with a Gaussian kernel (σ = 2 bins) at the single-trial level before averaging across trials. For each tertile, the resulting burst-rate time course was baseline-corrected by subtracting the subject-specific mean rate in the baseline window (-7 to -1 s), yielding a baseline-referenced rate time series (ΔHz) for each subject, hemisphere, and condition. Between-group differences in these baseline-corrected burst-rate time series were assessed using cluster-based permutation testing (two-sample; 1,000 permutations; two-tailed t-threshold based on the pooled degrees of freedom). Cluster significance was evaluated over the analysis interval (-7 to 14 s), and clusters shorter than 20 samples were discarded to reduce spurious detections.

Waveform-specific burst rates were further analyzed using linear mixed-effects models applied to window-averaged, baseline-referenced burst rates as a targeted follow-up to the time-resolved permutation tests, allowing us to quantify whether group differences reflected a consistent shift in burst rate within specific waveform-defined burst motifs. For each burst tertile, hemisphere, and execution condition (ExF, ExG), we computed a subject-level summary measure defined as the difference between mean burst rate during the task window (1-9 s) and the baseline window (-7 to -1 s). These values were then entered into mixed-effects models to assess between-group differences. Models were fit separately for each PC (PC6, PC8), tertile, hemisphere (contralateral, ipsilateral), and execution condition. In each model, burst rate was modeled as a function of group (DCD, comparison), with participant included as a random intercept to account for repeated measures. Statistical significance of fixed effects was assessed using Type III Wald χ² tests (car v3.1.0; Fox & Weisberg, 2018). Model fitting was performed in R (v4.2.2; R Core Team, 2022) using the lme4 package (v1.1.30; Bates et al., 2014).

### 2.9 Relationship between waveform-specific burst rates and motor skill (MABC-2)

To test whether execution-related, waveform-specific beta burst dynamics were associated with motor skill, we related baseline-referenced burst rates to MABC-2 total scores assessed following the EEG acquisition. Analyses were performed on ipsilateral sensorimotor electrode burst-rate time courses from the execution condition and PC identified in the group-difference analyses (PC6 during ExG; PC8 during ExF). For each PC, bursts were stratified into tertiles based on the global percentiles of PC score across all bursts. Within each participant and tertile, burst rates were computed by binning burst peak times in 200 ms intervals, converting counts to rates (Hz), smoothing at the single-trial level (Gaussian kernel; σ = 2 bins), averaging across trials, and baseline-referencing by subtracting the mean rate in the baseline window (-7 to -1 s). A single summary value per participant was then obtained as the mean baseline-referenced burst rate during the early execution period (1 to 6 s).

Associations between burst-rate summaries and MABC-2 scores were assessed using Spearman rank correlations pooling participants across groups and including participants from the comparison group with MABC-2 scores below the 25th percentile. To visualize and complement the rank-based effects, we fit separate group-wise linear models and tested the slope using OLS with HC3 heteroskedasticity-consistent standard errors. For slopes with *p* < 0.05, we plotted regression lines with 95% bootstrap confidence intervals (10,000 iterations; resampling participants with replacement). As a sensitivity analysis, we repeated the pooled correlation analyses using partial Spearman rank correlations controlling for age and IQ to determine whether the observed brain-behaviour relations were independent of developmental and cognitive factors.

### 2.10 Granger Causality Analysis

To test whether group differences in waveform-specific burst dynamics were accompanied by altered directed inter-hemispheric interactions, we estimated Granger causality (GC) from motif-specific burst-count time series rather than from the continuous beta envelope. This choice was motivated by the aim of quantifying directed coupling specifically between discrete burst events stratified by waveform motif (PC-score bins), whereas beta-envelope signals pool across motifs and can conflate changes in burst probability with changes in within-burst amplitude.

GC was computed only for the PC-condition pairs that showed robust motif-specific group effects in the burst-rate time series (PC6 during ExG; PC8 during ExF). For each analysis (PC6-ExG; PC8-ExF), bursts were first stratified into PC tertiles (T1-T3) using global PC-score percentiles. Within each subject, bursts were then converted into trial-wise point-process count series by binning burst peak times into 10 ms bins over the movement period (0-10 s post-onset), separately for ipsilateral and contralateral sensorimotor electrodes. This yielded six units per subject (ipsi T1-T3; contra T1-T3), each represented as an array of shape *units × trials × time bins*.

Directed connectivity was estimated using a point-process GC framework implemented as a negative-binomial GLM with constant dispersion (α = 1.0), following the logic of point-process Granger methods (Agouram et al., 2025; Kim et al., 2011). For each source→target unit pair, we fit (i) a full model predicting the target’s binned counts from an intercept, the lagged activity of the source, and the lagged activity of all other units (fully conditional GC), and (ii) a reduced model identical to the full model but omitting the source term. GC strength was defined as the likelihood-ratio statistic, GC = 2(LL_full_ - LL_reduced_). To aid interpretation, GC values were signed according to the sign of the source coefficient in the full model, with positive values indicating a positive directed influence and negative values indicating a negative directed influence in the point-process model. Because the burst detector prevents temporally overlapping bursts within a channel, within-hemisphere (and within-unit) dependencies are trivially constrained; therefore, within-hemisphere connections were excluded a priori and inference focused on inter-hemispheric connections.

History lags were selected per source-target pair by cross-validation over trials, choosing the lag that maximized held-out predictive performance. Candidate lags were 1-9 bins, corresponding to 10-90 ms given the 10 ms bin size, and model selection used 3-fold cross-validation. Statistical significance of within-group GC was assessed non-parametrically using 1,000 permutations. For each permutation and subject, trial labels were shuffled to disrupt across-unit temporal alignment for off-diagonal (inter-unit) interactions, while within-trial time points were shuffled to generate null values for diagonal terms; null GC matrices were then averaged across subjects to form a group-level null distribution for each connection. Two-sided p-values were computed as the proportion of permuted GC values with absolute magnitude exceeding the observed absolute GC value, and multiple comparisons across connections were controlled using Benjamini-Hochberg FDR (α = 0.05) within each PC-condition analysis. Between-group differences were tested by subject-label permutation (10,000 permutations), recomputing the group-mean signed-GC difference (DCD - comparison) for each connection and deriving two-sided p-values from the resulting null distribution; significant group-difference connections were retained for visualization and interpretation after the same within-analysis multiple-comparisons control.

Source code for all of these analyses is available at https://github.com/danclab/dcd_bursts.

## 3. Results

### 3.1 Time-frequency-based burst features do not differ between DCD and comparison groups

We first examined conventional beta power dynamics over sensorimotor cortex (C3/C4), baseline-corrected to the pre-task interval. Cluster-based permutation testing of the continuous time series revealed task-related beta modulation only during motor execution (Figure 3, see Figure S1 for observation conditions). During fine motor execution (ExF), significant contralateral beta modulation was observed in the comparison group, whereas no reliable clusters were detected in the DCD group. During gross motor execution (ExG), significant beta modulation was present ipsilaterally in both groups. No significant between-group differences emerged in any condition from the time-resolved analyses. Importantly, task epochs lasted 10 s and were not time-locked to individual movement events, limiting the temporal specificity of conventional power estimates and motivating a complementary window-based analysis.

**Figure 3.**
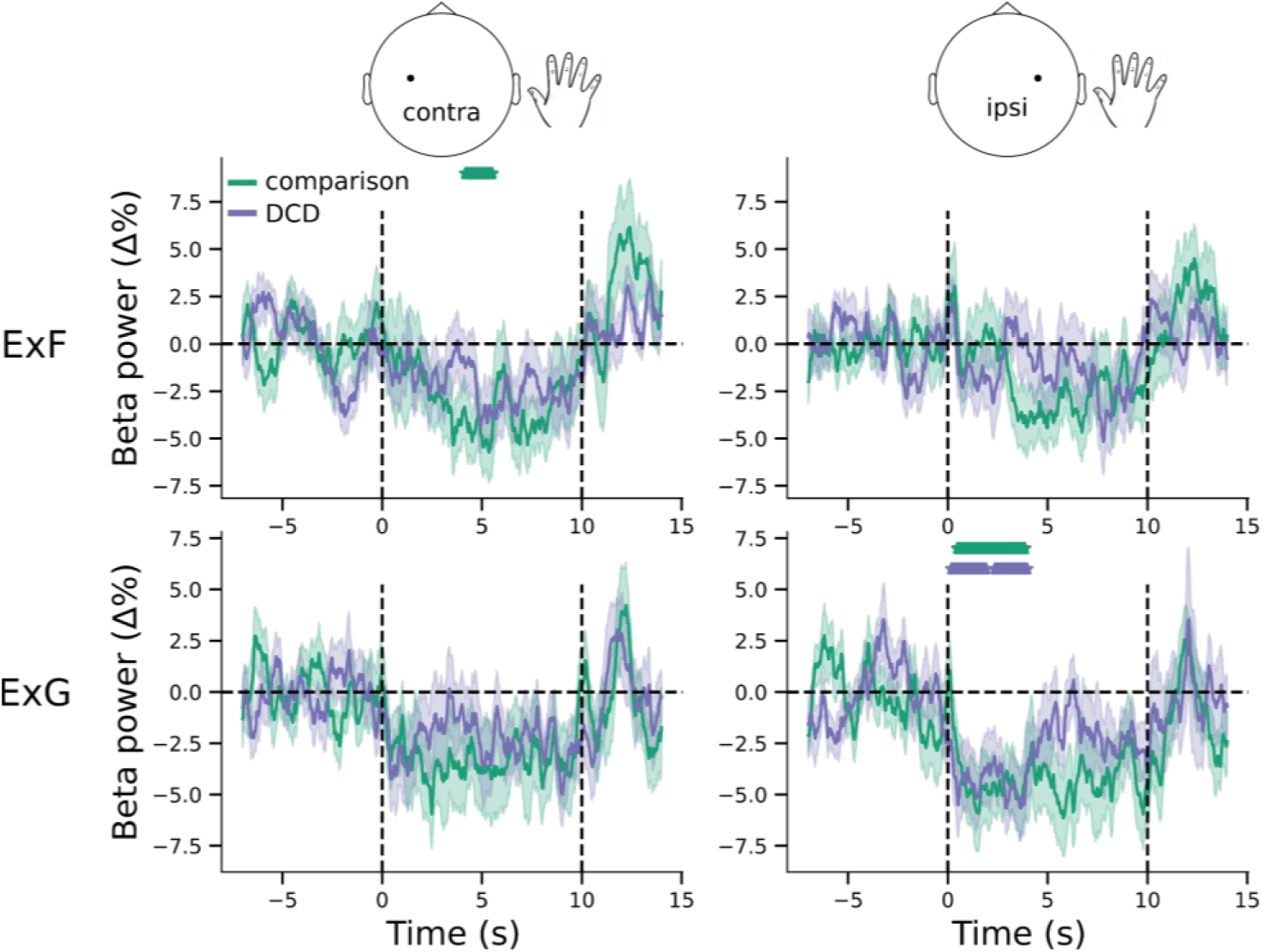
Task-related modulation of sensorimotor beta power during fine and gross motor execution. Time courses of baseline-corrected beta power (% change from baseline) over sensorimotor cortex during execution of fine (ExF) and gross (ExG) actions, shown separately for the hemisphere contralateral (left column) and ipsilateral (right column) to the hand used. Solid lines depict group means for the comparison (green) and DCD (purple) groups; shaded regions indicate ±SEM across participants. Vertical dashed lines mark task onset (0 s) and offset (10 s); the horizontal dashed line indicates baseline (0%). Colored asterisks denote significant within-group deviations from baseline (one-sample permutation tests, *p* < 0.05).

We therefore averaged beta power within three predefined windows (baseline, task, and post-task) and analysed trial-wise values using linear mixed-effects models. During contralateral ExF, there was a significant main effect of window (*χ*²(2) = 20.77, *p* < 0.001) and a group × window interaction (*χ*²(2) = 7.26, *p* = 0.026), but no main effect of group (*χ*²(1) = 1.87, *p* = 0.171). Post hoc contrasts indicated reduced beta power during the task relative to baseline in both groups (both *p* ≤ 0.002), consistent with movement-related beta desynchronization, with a larger reduction in the comparison group; however, no group differences were observed within any individual window (all *p* > 0.10), including the baseline period (*t*(36) = -1.37, *p* = 0.180). Ipsilateral ExF showed a similar interaction (*χ*²(2) = 6.29, *p* = 0.043) without a main effect of group (*χ*²(1) = 1.87, *p* = 0.171), with no differences in baseline beta power (*t*(35.8) = -1.37, *p* = 0.180).

For ExG, both hemispheres exhibited a robust main effect of window (contralateral: *χ*²(2) = 12.46, *p* = 0.002; ipsilateral: *χ*²(2) = 15.23, *p* < 0.001), again reflecting reduced beta power during task relative to baseline (all *p* ≤ 0.0004), but neither hemisphere showed a main effect of group (both *p* ≥ 0.94) nor a group × window interaction (both *p* ≥ 0.23). Accordingly, no evidence for group differences in baseline beta power was observed during ExG. Importantly, there was no consistent increase in post-task beta power relative to baseline in any condition, indicating the absence of a reliable beta rebound at the group level. These findings indicate that conventional beta power captures expected movement-related desynchronization but does not reveal stable group differences. Moreover, the absence of group differences in baseline beta power indicates that the relative power effects cannot be attributed to baseline differences or ceiling/floor effects.

We next assessed whether sensorimotor beta activity exhibited sustained rhythmic structure using lagged Hilbert autocoherence (LHaC) (Zhang et al., 2025). Across groups, LHaC values were lower within the identified beta range (Figure 4a-b) relative to lower frequencies, indicating limited long-range periodicity and supporting the characterization of beta activity as transient rather than sustained rhythmic oscillations (Figure 4c-d). No qualitative differences between groups were apparent in the spectral or autocoherence profiles.

**Figure 4.**
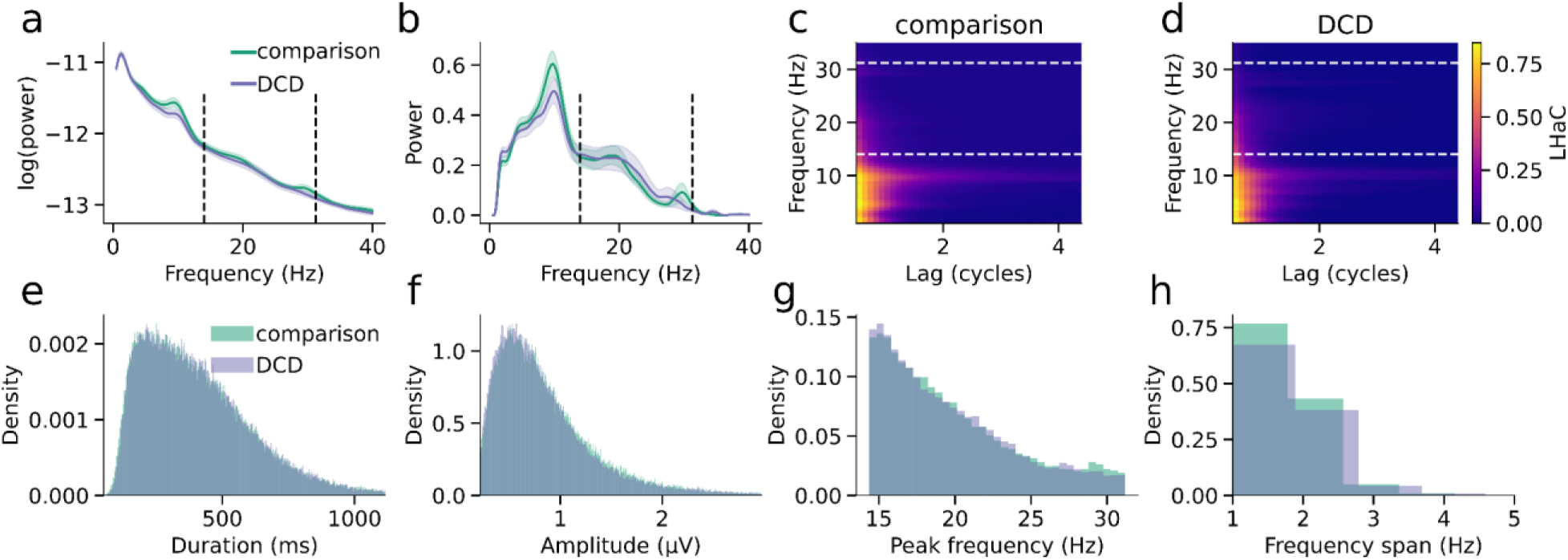
Spectral characteristics and time-frequency burst features do not differ between groups. (a) Log power spectral density over central sensorimotor electrodes (C3/C4), averaged across participants, for comparison (green) and DCD (purple) groups; shaded areas indicate ±SEM. (b) Periodic (aperiodic-corrected) power spectra derived using spectral parameterization. Vertical dashed lines in (a-b) mark the beta band used for burst detection (14-31.25 Hz). (c-d) Mean lagged Hilbert autocoherence (LHaC) as a function of frequency (≤35 Hz) and lag (cycles), averaged across hemispheres, shown for the comparison group (c) and DCD group (d); white dashed lines indicate the beta band. Color scale is matched across groups. (e-h) Distributions of burst features pooled across task conditions: duration (FWHM in time; ms) (e), peak amplitude (µV) (f), peak frequency (Hz) (g), and frequency span (FWHM in frequency; Hz) (h). Across all spectral and burst-level metrics, groups did not differ significantly.

We then applied an iterative burst detection algorithm (Rayson et al., 2023; Szul et al., 2023) to identify candidate beta burst events and compared time-frequency-based burst features between groups, pooling bursts detected over left and right sensorimotor cortex (C3/C4). There were no group differences in burst duration (FWHM time; *M_comparison_* = 0.420 s, *SD* = 0.049; *M_DCD_* = 0.418 s, *SD* = 0.053; *p* = 0.868), peak amplitude (*M_comparison_* = 0.943 μV, *SD* = 0.321; *M_DCD_* = 0.940 μV, *SD* = 0.500; *p* = 0.979), peak frequency (*M_comparison_* = 19.54 Hz, *SD* = 1.17; *M_DCD_* = 19.66 Hz, *SD* = 1.39; *p* = 0.762), or frequency span (FWHM frequency; *M_comparison_* = 1.46 Hz, *SD* = 0.09; *M_DCD_* = 1.45 Hz, *SD* = 0.13; *p* = 0.964; Figure 4e-h). Finally, we examined overall burst rate across execution (ExF, ExG) conditions, expressed relative to the pre-task baseline. Burst rate did not differ between groups in any condition (all *p* < 0.152). Conventional time-frequency burst metrics and burst incidence therefore do not distinguish DCD from comparison participants, motivating a more detailed analysis of burst waveform structure.

### 3.2 Sensorimotor cortical bursts share similar waveform motifs across groups

Because time-frequency-based beta burst features are derived from time-frequency decomposition of the burst event time series, and the correspondence between temporal and time-frequency-based burst features is not one-to-one (Jones, 2016; Szul et al., 2023), we then focused our analysis on burst waveform shapes. In line with previous work, we found that sensorimotor cortical bursts in both groups had a wavelet-like median waveform shape with a strong central negative deflection and surrounding positive deflections (Bonaiuto et al., 2021; Brady & Bardouille, 2022; Rayson et al., 2023; Figure 5a; Sherman et al., 2016; Szul et al., 2023). To assess group differences in burst waveform shape, we next applied PCA to the burst waveforms, an approach previously shown to capture functionally relevant waveform motifs (Rayson et al., 2023; Szul et al., 2023). We ran separate PCAs on burst waveforms from each group and compared the resulting eigenvectors. All PCs were highly similar between groups (*ρ* = 0.85 - 1.0; Figure 5b). For nearly all PCs, the most similar component from the comparison group PCA was the corresponding one of the DCD group PCA, indicating that the waveform motif and relative percentage of variance explained was very similar across groups.

**Figure 5.**
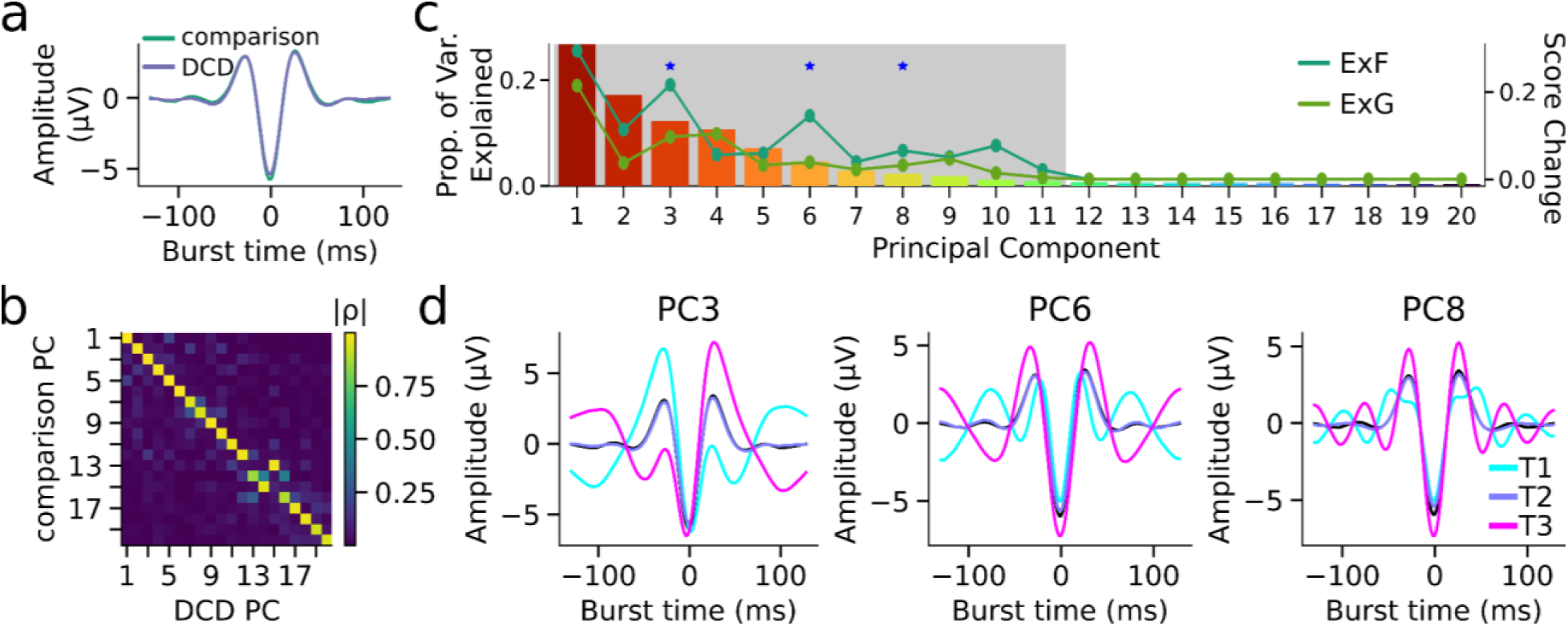
Burst waveform motifs and task-related modulation revealed by PCA. (a) Median burst waveforms over sensorimotor cortex for comparison (green) and DCD (purple) groups, showing highly similar canonical beta burst morphology. (b) Absolute Spearman correlation coefficients between eigenvectors obtained from separate PCAs performed on comparison and DCD bursts, demonstrating strong correspondence of waveform components across groups. (c) Proportion of variance explained by each principal component (bars). Grey shading indicates components exceeding the permutation-based significance threshold derived from waveform shuffling. Overlaid lines show the mean absolute change in PC score from baseline during execution of fine and gross actions (averaged across groups). Blue asterisks mark components exhibiting significant execution-related modulation. (d) For each significantly modulated component, mean waveform shapes of bursts falling within each score tertile (T1-T3; colored lines), plotted against the overall mean waveform (black), illustrating distinct waveform motifs captured along each principal axis.

### 3.3 Specific burst waveform motifs are modulated by action execution tasks

Given the similarity in waveform shape and variability between comparison and DCD bursts, we next performed a single global PCA on all sensorimotor burst waveforms to represent bursts within a common low-dimensional space. Components retained for further analysis were determined using a temporal-shuffling permutation procedure, which identified those PCs whose variance exceeded that expected under waveform phase randomization (PCs 1-11, all *p* < 0.001). For each retained component, we then quantified condition-related modulation by computing, separately for each group, the difference in mean PC score between execution (ExF, ExG) and pre-task baseline, averaging within subjects and testing significance using paired permutation tests. This analysis revealed selective modulation during gross motor execution (ExG) in the comparison group only. Specifically, PC3 (*p* = 0.019), PC6 (*p* = 0.042), and PC8 (*p* = 0.045) showed significant baseline-to-task changes in the comparison group. No PCs showed significant modulation during fine motor execution (ExF), and no components reached significance in the DCD group for either condition. Each of these components captures distinct deviations from the mean burst waveform, reflecting variation in the depth of the central negative deflection and the relative prominence of surrounding peaks. Because changes in mean PC score are ambiguous with respect to burst incidence (i.e., an increase in mean score could reflect increased occurrence of high-scoring bursts or decreased occurrence of low-scoring bursts), we next examined these components in terms of waveform-specific burst rate.

### 3.4 Waveform-specific burst rates differ between DCD and comparison groups

For each PC that showed task-related modulation, we quantified waveform-specific burst rate by binning bursts into tertiles of PC score and computing the mean burst rate (Hz) within three predefined epoch windows (baseline, task, and post-task). Burst-rate time courses were estimated separately for contralateral and ipsilateral sensorimotor electrodes during ExF and ExG. Cluster-based permutation testing of the burst-rate time series revealed ipsilateral group differences that were selective to bursts with specific waveform shapes, with significant between-group effects for the first and third tertiles of PC6 during ExG (Figure 6) and the third tertile of PC8 during ExF (Figure 7); no other PCs or hemispheres showed reliable task-related group differences.

**Figure 6.**
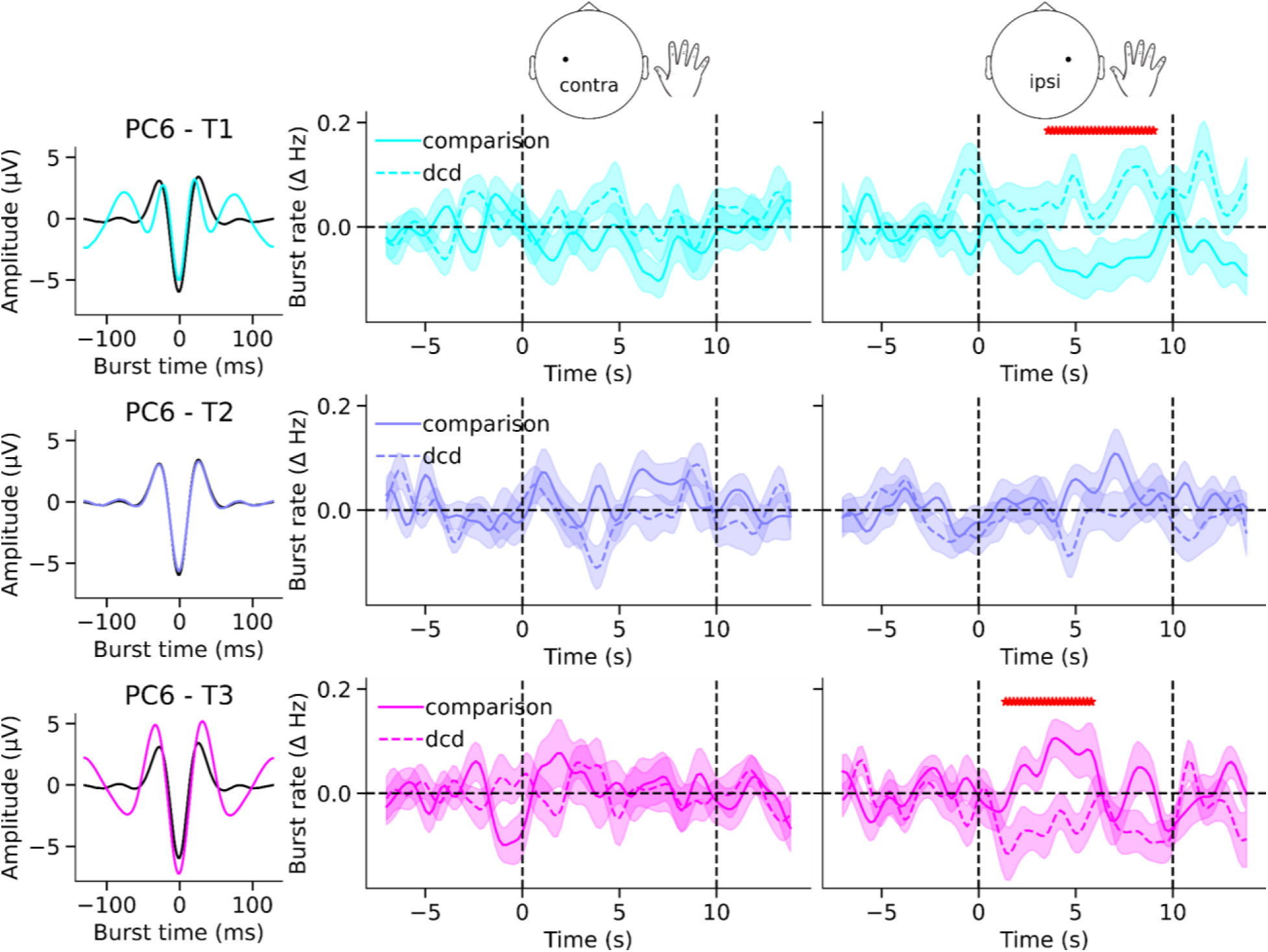
Waveform-specific burst-rate differences between groups along PC6 during gross motor execution (ExG). Left column: Mean waveform shapes of bursts falling into each tertile (T1-T3; color-coded from cyan to magenta) of PC6 scores, shown relative to the overall mean waveform (black), illustrating the distinct waveform motifs captured along this principal axis. Middle and right panels: Time-resolved, baseline-corrected burst rates (ΔHz) for comparison (solid lines) and DCD (dashed lines) groups during gross motor execution (ExG), plotted separately for the hemisphere contralateral and ipsilateral to the dominant hand. Shaded regions indicate ±SEM across participants. Vertical dashed lines mark task onset (0 s) and offset (10 s); the horizontal dashed line indicates baseline (0 Hz). Red asterisks denote significant between-group clusters identified using cluster-based permutation testing (*p* < 0.05).

**Figure 7.**
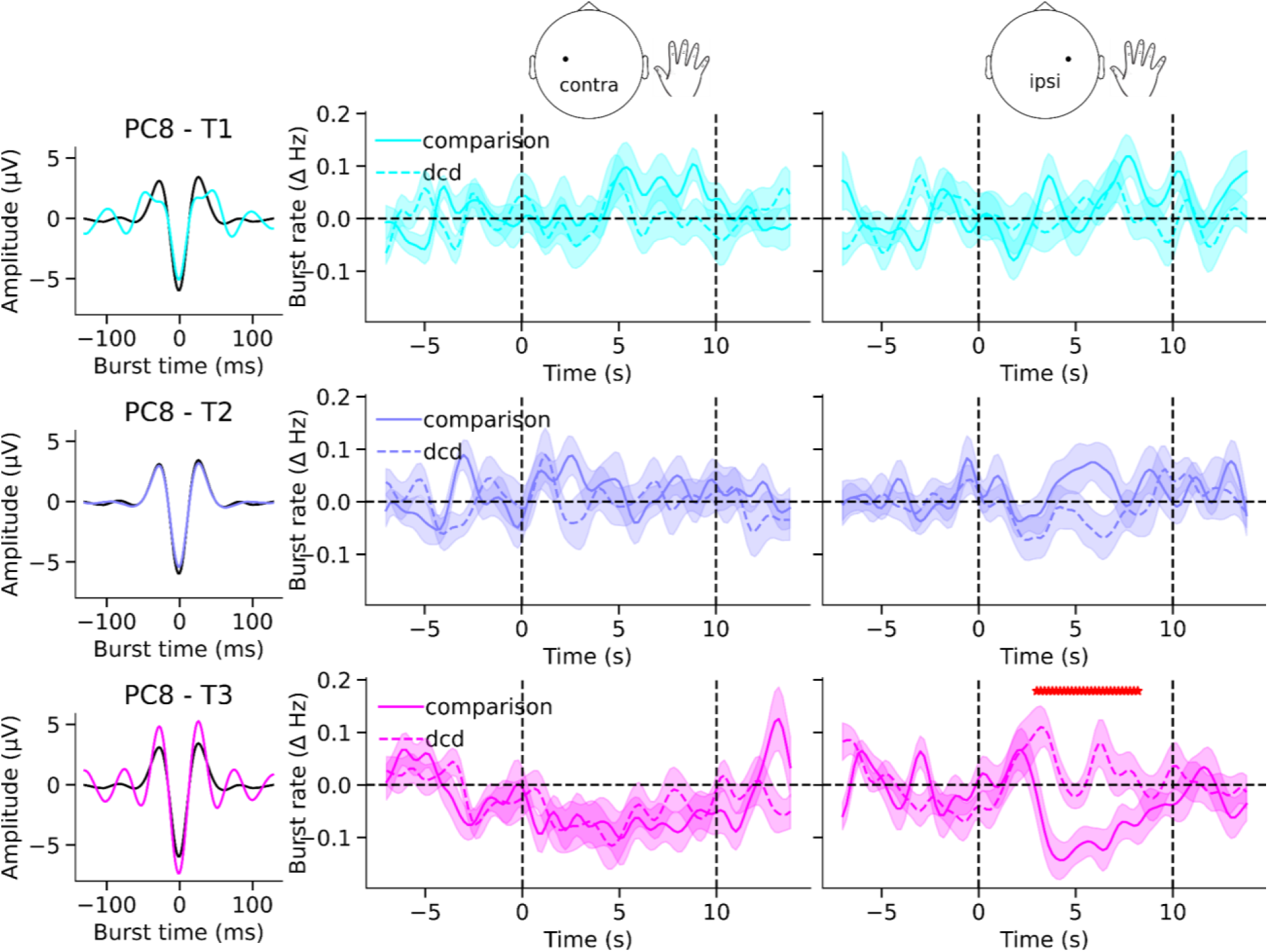
Waveform-specific burst-rate differences between groups along PC8 during fine motor execution (ExF). Left column: Mean waveform shapes of bursts falling into each tertile (T1-T3; color-coded from cyan to magenta) of PC8 scores, shown relative to the overall mean waveform (black), illustrating the distinct waveform motifs captured along this principal axis. Middle and right panels: Time-resolved, baseline-corrected burst rates (ΔHz) for comparison (solid lines) and DCD (dashed lines) groups during fine motor execution (ExF), plotted separately for the hemisphere contralateral and ipsilateral to the dominant hand. Shaded regions indicate ±SEM across participants. Vertical dashed lines mark task onset (0 s) and offset (10 s); the horizontal dashed line indicates baseline (0 Hz). Red asterisks denote significant between-group clusters identified using cluster-based permutation testing (*p* < 0.05).

Importantly, these effects reflected opposing task-related modulations across the extremes of the PC dimensions rather than uniform changes in overall bursting. Cluster-based permutation tests of the time-resolved burst rate revealed cross-over patterns in the ipsilateral hemisphere during movement execution. For PC6 (ExG, ipsilateral), the lowest-score tertile (T1) showed a decrease in burst rate during movement in the comparison group but an increase in the DCD group, whereas the highest-score tertile (T3) showed the inverse pattern (Figure 6). A similar divergence was observed for PC8 (ExF, ipsilateral), where the highest-score tertile (T3) exhibited a task-related decrease in burst rate in the comparison group and a concurrent increase in the DCD group (Figure 7). These patterns indicate a redistribution of burst occurrence toward opposite ends of the waveform dimensions between groups.

To quantify these effects, we performed follow-up linear mixed-effects analyses on window-averaged burst rates (task minus baseline), focusing on PCs and conditions identified in the time-resolved analysis. For PC6 during gross motor execution (ExG), significant between-group differences were confined to the ipsilateral hemisphere and were selective to the extreme tertiles. In the lowest-score tertile (T1), burst rate differed significantly between groups (*χ*²(1) = 8.59, *p* = 0.003), with the DCD group showing a higher task-related increase relative to the comparison group. In contrast, the highest-score tertile (T3) showed the opposite pattern (*χ*²(1) = 5.55, *p* = 0.018), reflecting a reduction in burst rate in the DCD group relative to controls. No group effects were observed for the middle tertile (T2) or in the contralateral hemisphere (all *p* > 0.12). For PC8 during fine motor execution (ExF), a significant group effect was observed only for the ipsilateral highest-score tertile (T3) (*χ*²(1) = 8.96, *p* = 0.0028). Here, the comparison group exhibited a task-related decrease in burst rate, whereas the DCD group showed a relative increase. No significant group effects were present for lower tertiles, the contralateral hemisphere, or during ExG (all *p* > 0.13). Together, these analyses converge to show that execution-related group differences are waveform-specific, ipsilaterally expressed, and restricted to the extremes of particular waveform dimensions (PC6 and PC8). Rather than reflecting a uniform shift in burst rate, these effects indicate a redistribution of burst occurrence across distinct waveform motifs in DCD, consistent with altered engagement of specific beta burst subtypes during motor execution.

### 3.5 Waveform-specific burst rates along PC6 and PC8 correlate with motor skill

Given that execution-related group differences were selective to ipsilateral burst subtypes defined by PC6 (ExG) and PC8 (ExF), we next asked whether these task-related, baseline-corrected burst-rate changes scaled with motor performance more generally (Figure 8). To capture variability across the full sample, we included all participants, including comparison-group children with MABC-2 scores below the 25th percentile. We quantified, for each participant, the mean task-baseline change in burst rate for bursts in each tertile of PC score (ipsilateral hemisphere; ExG for PC6 and ExF for PC8), and related these values to MABC-2 total score using Spearman rank correlations.

**Figure 8.**
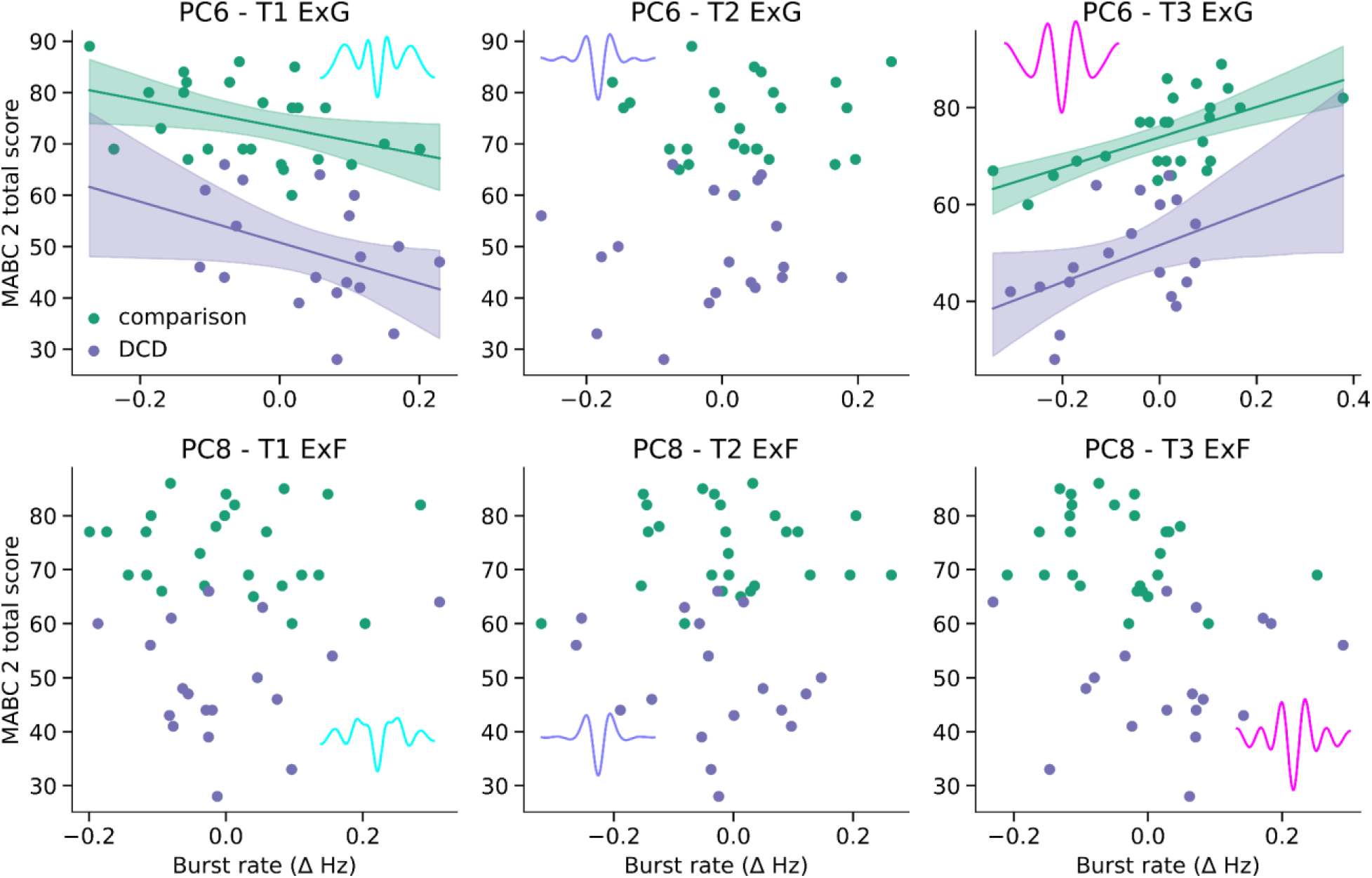
Waveform-specific burst rates predict motor performance in a motif- and condition-dependent manner. Scatter plots show the relationship between baseline-corrected burst rate (ipsilateral hemisphere) and MABC-2 total score for bursts falling into each tertile (T1-T3) of the indicated principal component (PC6, PC8) during execution conditions. Points represent individual participants (comparison: green; DCD: purple). Solid lines indicate group-wise linear fits, with shaded regions denoting bootstrapped 95% confidence intervals (10,000 iterations); fits are shown where *p* < 0.05.

For PC6 during ExG, motor performance was strongly related to the two extreme tertiles in opposite directions: T1 burst-rate change was negatively associated with MABC-2 (*ρ* = -0.55, *p* < 0.001), whereas T3 burst-rate change was positively associated with MABC-2 (*ρ* = 0.54, *p* < 0.001). Robust (HC3) group-wise linear fits were consistent with this pattern: for PC6-T1, slopes were negative in both groups (comparison: slope = - 28.85, *p* = 0.020; DCD: slope = -47.03, *p* = 0.039), and for PC6-T3, slopes were positive in both groups (comparison: slope = 30.63, *p* < 0.001; DCD: slope = 38.07, *p* = 0.023). This mirrors the redistribution observed in the group contrasts: better motor performance corresponded to fewer low-PC6 (T1) bursts and more high-PC6 (T3) bursts during ExG in both groups. For PC8 during ExF, the association with motor performance was specific to the highest tertile (T3), where task-baseline burst-rate change was negatively correlated with MABC-2 (*ρ* = −0.42, *p* = 0.004). However, this relationship was not reliable within either group alone using robust linear fits (comparison: slope = -23.52, *p* = 0.099; DCD: slope = 9.22, *p* = 0.760), suggesting that the effect is expressed primarily across the pooled sample rather than as a consistent within-group slope. To determine whether the relation between waveform-specific burst rates and motor skill could be explained by developmental or cognitive factors, we repeated the analyses controlling for age and IQ. The principal associations remained significant after adjustment, including PC6 T1 (*ρ* = - 0.50, *p* < 0.001), PC6 T3 (*ρ* = 0.54, *p* < 0.001), and PC8 T3 (*ρ* = -0.30, *p* = 0.048), indicating that the observed brain-behaviour relations cannot be explained solely by age-or IQ-related differences.

### 3.6 Waveform-specific interhemispheric connectivity differences in DCD during motor execution

To test whether motif- and hemisphere-specific differences in burst dynamics were accompanied by altered directed interactions between sensorimotor cortices, we examined signed Granger causality (GC) between ipsilateral and contralateral burst-rate time series. GC was computed separately for each waveform tertile within a given PC and execution condition. Within-hemisphere interactions were excluded because the burst detection procedure precludes temporal overlap and therefore trivially constrains intra-hemispheric coupling. Statistical significance was assessed using permutation testing with FDR correction for within-group effects and subject-label permutation for between-group contrasts.

For PC6 during gross motor execution (ExG), clear group differences in interhemispheric connectivity were observed (Figure 9a-c). Two directed connections differed significantly between groups. Connectivity from ipsilateral T1 to contralateral T2 showed a negative directed influence in the comparison group but a positive directed influence in the DCD group, indicating a reversal in the sign of directional influence. In addition, connectivity from contralateral T3 to ipsilateral T1 showed a positive directed influence in the comparison group but a negative directed influence in DCD, again reflecting a qualitative change in coupling rather than a simple attenuation. These effects show that, for PC6, DCD is associated with sign-inverted interhemispheric interactions between bursts at opposing ends of the waveform dimension. For PC8 during fine motor execution (ExF), group differences were more circumscribed (Figure 9d-f). A significant effect was observed for the connection from ipsilateral T1 to contralateral T3, which showed a significant negative directed influence in DCD but was not significant in the comparison group. No additional interhemispheric connections showed reliable group differences for PC8. Overall, the connectivity analyses reveal that group differences in beta burst dynamics are accompanied by selective disruptions of interhemispheric communication, expressed in a waveform-dependent manner and, in some cases, involving sign-inverted interhemispheric interactions during motor execution.

**Figure 9.**
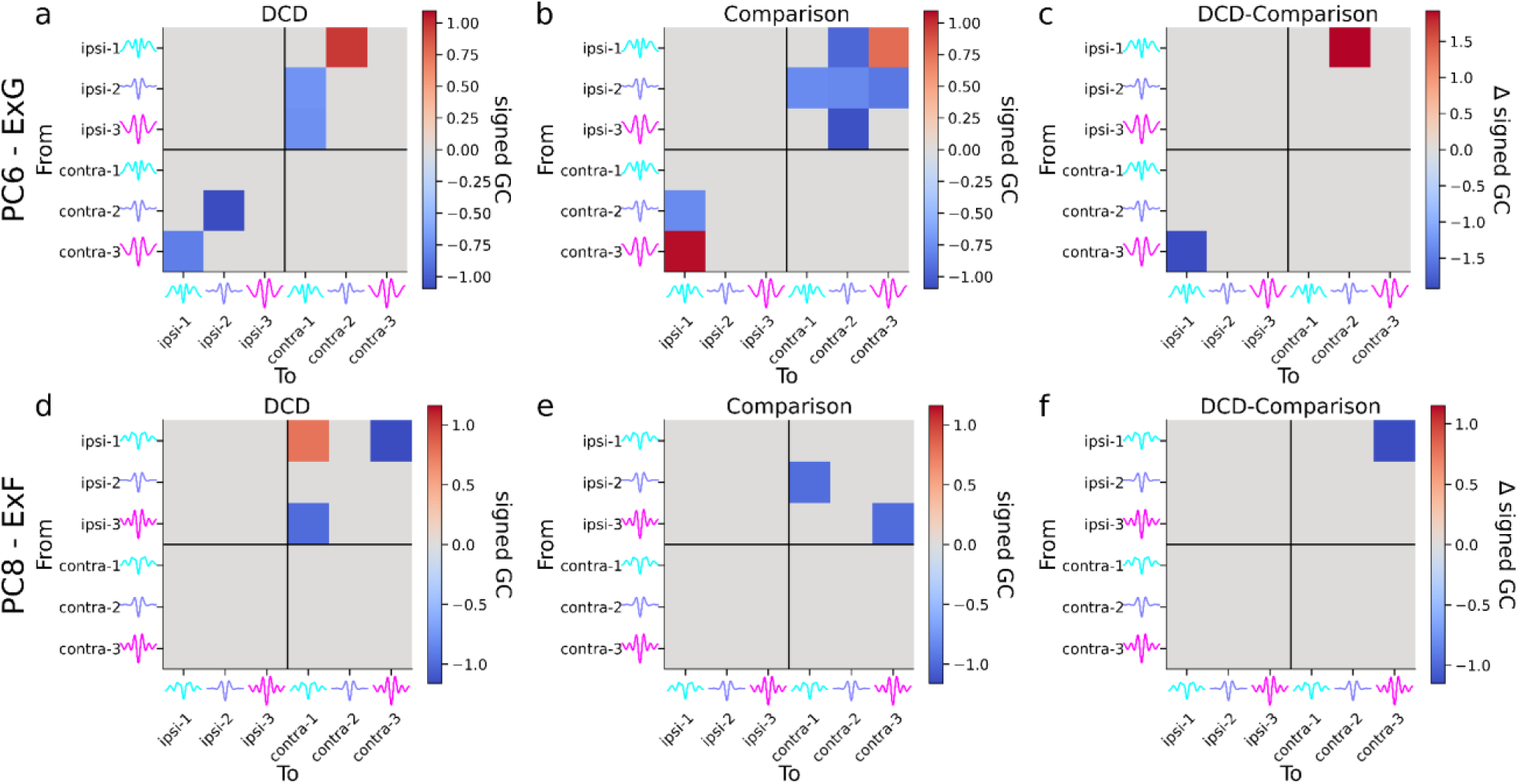
Waveform-specific, hemisphere-resolved Granger causality during execution for PC6 (ExG) and PC8 (ExF). Signed Granger causality (GC) matrices computed from burst-rate time series (0-10 s post-movement onset) for bursts stratified by waveform tertile within a given principal component. Units correspond to ipsilateral and contralateral sensorimotor electrodes, separated by black lines; within each hemisphere, rows and columns represent PC tertiles (T1-T3; waveform motifs illustrated). Top row (PC6, ExG): (a) DCD group; (b) comparison group; (c) between-group difference (DCD - comparison). Bottom row (PC8, ExF): (d) DCD group; (e) comparison group; (f) between-group difference (DCD - comparison). Colored cells indicate significant GC values after permutation testing with FDR correction (within-group panels) or significant group differences from subject-label permutation (right panels); non-significant connections are set to zero. Warmer colors denote stronger positive signed GC (directional influence), and cooler colors denote negative signed GC.

## 4. Discussion

This study applied waveform-based beta burst analyses to investigate how sensorimotor burst motifs relate to motor performance in childhood. Conventional time-frequency measures, including beta power and burst spectral features, did not differ between groups, and the overall structure of burst waveform variability was highly similar in children with and without developmental coordination disorder. Instead, differences emerged only when examining burst dynamics along specific waveform dimensions. Movement execution was associated with systematic redistribution of bursts along these dimensions, and the prevalence and lateralization of particular motifs co-varied with motor coordination. Children with DCD were disproportionately represented at one end of this motif-defined axis, but the relation between waveform-specific burst rate and MABC-2 scores followed a similar slope across groups. These findings indicate that beta bursts comprise functionally heterogeneous events (Moreau et al., 2026; Rayson et al., 2023, 2025; Szul et al., 2023; Zich et al., 2023) whose motif-specific dynamics track continuous variation in motor competence, with DCD providing a clinically informative model of extreme motor difficulty rather than a categorically distinct beta phenotype.

Our results further suggest that different beta burst waveform motifs contribute differently to sensorimotor processing during action execution. Rather than reflecting discrete burst classes, the principal components derived from waveform analysis capture continuous dimensions of burst shape variability along which bursts can be redistributed depending on task demands and motor performance. Group differences emerged selectively at the extremes of these dimensions: during gross motor execution, bursts with low versus high PC6 scores showed opposing changes in rate across groups, while during fine motor execution a similar divergence was observed for bursts occupying the highest tertile of PC8. Importantly, these effects did not involve global changes in burst incidence or spectral burst features. Instead, motor-related differences reflected shifts in the relative prevalence of bursts occupying specific regions of the waveform variability space. This pattern suggests that sensorimotor beta bursts form a heterogeneous population of transient events whose functional contribution depends on their waveform motif.

A notable feature of these effects was their localization to the ipsilateral hemisphere. Although sensorimotor beta activity is often interpreted primarily in terms of contralateral motor control, ipsilateral sensorimotor cortex plays an important role in coordinating activity across hemispheres during movement (Serrien et al., 2006). The motif-specific redistribution observed here may therefore reflect differences in how transient beta events participate in interhemispheric sensorimotor communication (Seedat et al., 2020). Consistent with this interpretation, directed connectivity analyses revealed motif-specific differences in interhemispheric interactions, including changes in the sign of influence for particular burst tertiles. These findings suggest that the coordination of activity between hemispheres depends not only on the timing of beta bursts but also on the waveform motifs through which these events are expressed.

Although motif-specific differences in directed interhemispheric interactions were associated with differences in burst waveform expression, the present analyses do not establish the direction of this relationship. Altered recruitment of specific burst motifs may influence coordination between bilateral sensorimotor networks, but it is equally plausible that changes in network coordination shape the expression of particular burst motifs, or that both arise from a common underlying mechanism. Distinguishing between these possibilities will require future studies combining perturbation approaches, computational modelling, or task manipulations that explicitly probe interhemispheric coordination.

The relation between burst dynamics and motor performance further supports a dimensional interpretation at the level of brain-behavior mapping. Although children with DCD exhibited poorer motor performance on the MABC-2 overall, the association between waveform-specific burst rates during the EEG execution conditions and MABC-2 score was comparable across groups, indicating that similar neural-behavioral relationships are expressed as a continuum across the full sample, rather than reflecting diagnostic status alone. Moreover, these associations remained significant after controlling for age and IQ, suggesting that they are not simply driven by developmental or cognitive differences between participants. This pattern suggests that variation in sensorimotor beta burst dynamics may index sensorimotor performance, rather than an underlying process specific to DCD. Although DCD is defined using cut-off criteria on standardized motor assessments, motor skill is continuously distributed across the population. The overlap in MABC-2 scores between groups in the present sample suggests that waveform-specific beta burst dynamics may track individual differences in motor skill across this continuum rather than diagnostic status alone. This dimensional interpretation may help explain the significant associations between burst dynamics and motor skill despite the absence of robust categorical group differences across most conventional EEG measures.

The present findings therefore complement existing accounts of motor impairment in DCD that emphasize atypical sensorimotor connectivity and hemispheric specialization (McLeod et al., 2014, 2016; Tallet et al., 2013). Although previous work has linked these differences to alterations in corpus callosum-mediated motor coordination and interhemispheric inhibition, the present unimanual visuomotor control task did not directly assess these mechanisms. Consequently, our findings should be interpreted as evidence for altered functional interactions between bilateral sensorimotor networks during movement execution, rather than as evidence for altered transcallosal inhibitory processing. The motif-specific redistribution observed here provides a potential physiological substrate for these network-level differences. If distinct beta burst motifs reflect different modes of sensorimotor processing, then altered recruitment of these motifs could influence how information is exchanged between hemispheres during action planning and execution. Such differences may ultimately contribute to the motor planning and adaptation difficulties often observed in DCD (Subara-Zukic et al., 2022). More broadly, these findings suggest that the functional role of beta bursts in motor control may depend on the distribution of waveform motifs within the burst population rather than on overall burst incidence alone.

Task demands may also shape how these motif dynamics are expressed. The motif-specific differences observed here varied between fine and gross motor conditions, suggesting that the recruitment of particular burst motifs depends on the structure of the motor task. Fine motor actions require precise and sustained control of a specific effector, whereas gross motor actions often involve broader muscle synergies and more distributed sensorimotor engagement. Differences in effector specificity and bilateral coordination may therefore influence how burst motifs are distributed across hemispheres and how strongly the ipsilateral sensorimotor cortex contributes to task execution. In children with DCD, altered motif recruitment during these tasks may reflect differences in the ability to coordinate sensorimotor activity across hemispheres when task demands require flexible integration of bilateral motor signals, for example when stabilizing one hand while executing a precise action with the other, or when suppressing unintended mirror movements.

The present study has several strengths. The inclusion of execution, observation, and non-biological motion conditions allowed us to isolate sensorimotor-specific effects and demonstrate that motif-related differences were not simply driven by visual or attentional factors. Moreover, this study represents the first application of waveform-based beta burst analysis to a neurodevelopmental population, demonstrating the utility of this approach for characterizing subtle differences in sensorimotor network dynamics that are not apparent using conventional spectral measures. Nevertheless, several limitations should be noted. The sample size, while comparable to previous EEG studies in DCD, was relatively modest, and replication in larger cohorts will be important. We also did not quantify participants’ prior motor experience (e.g., participation in sports, musical training, or other structured motor activities), which may contribute to variability in motor performance and sensorimotor cortical dynamics independently of DCD diagnosis. Future studies should consider these experiential factors alongside neurodevelopmental measures. In addition, the present analyses focused on central electrodes overlying sensorimotor cortex; beta bursts arising from associative or prefrontal regions may also contribute to motor planning and control (Diesburg et al., 2021; Liljefors et al., 2024; Wessel, 2020). Because the experimental design used relatively long task epochs, the temporal relation between individual movements and burst events could not be examined with high precision. Future studies using event-locked designs will be important for understanding how motif-specific bursts relate to individual movement components and error signals. Finally, although extensive preprocessing and trial exclusion procedures were used to minimise movement- and muscle-related artefacts, individual differences in attention, task engagement, or movement strategies may nonetheless have contributed to variability in beta burst dynamics. Given the relatively modest sample size, these factors cannot be fully disentangled from neurophysiological differences and should be examined more directly in future studies.

In conclusion, our findings demonstrate that sensorimotor beta burst dynamics in childhood vary systematically along waveform-defined dimensions that relate to motor performance. Rather than reflecting global differences in beta activity, motor coordination differences were associated with motif-specific redistribution of bursts within the broader population of beta events. These results support emerging models in which the functional heterogeneity of beta bursts plays a central role in sensorimotor computation. By linking waveform-defined burst motifs to motor competence across development, this work provides new insight into the neural mechanisms underlying motor coordination differences and highlights the potential of burst-based approaches for studying atypical motor development.

## Data and Code Availability

All analysis code used in this study is publicly available at: https://github.com/danclab/dcd_bursts. The dataset supporting the findings of this study is available via the Open Science Framework (OSF) at: https://osf.io/vnup9/.

## Author Contributions

H.R.: Conceptualization, Methodology, Software, Formal analysis, Visualization, Writing – Original Draft, Writing – Review & Editing. S.A.G.: Conceptualization, Supervision, Writing – Original Draft, Writing – Review & Editing. J.K.: Writing – Review & Editing. C.R.G.J.: Writing – Review & Editing. C.P.: Writing – Review & Editing. J.J.B.: Conceptualization, Methodology, Software, Formal analysis, Visualization, Writing – Original Draft, Writing – Review & Editing, Supervision, Funding acquisition. R.E.V.: Conceptualization, Investigation, Data curation, Supervision, Writing – Review & Editing, Funding acquisition.

