## Supplementary Material for "Functional heterogeneity of beta bursts in childhood reveals a dimensional neural signature of motor skill"

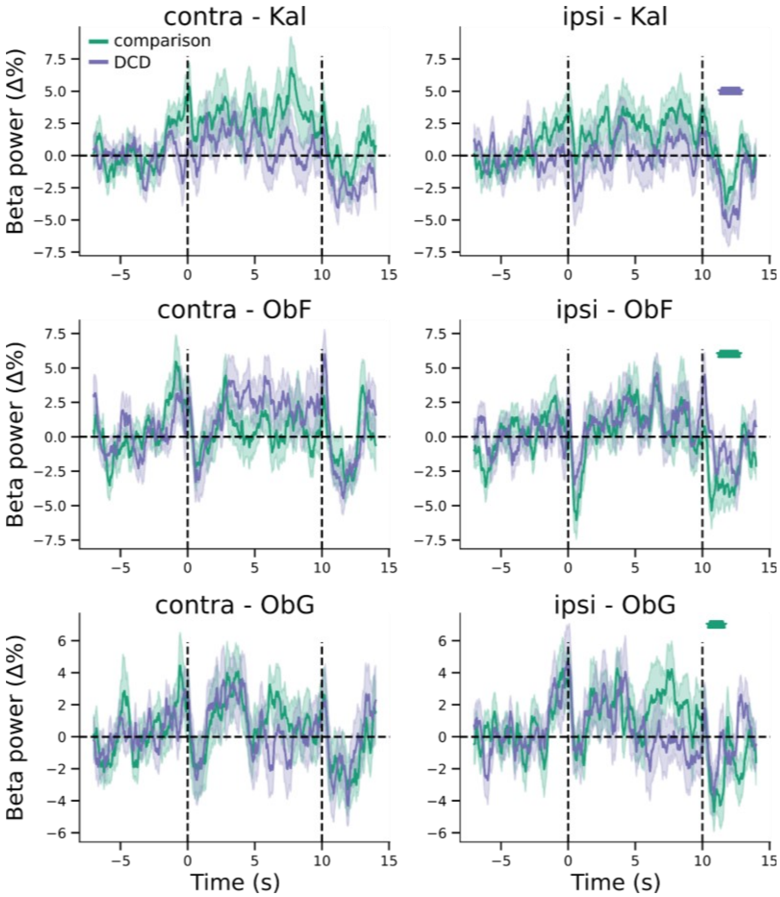


**Figure S1. Task-related modulation of sensorimotor beta power during observation conditions.** Time courses of baseline-corrected beta power (% change from baseline) over sensorimotor cortex during observation of a kaleidoscope (Kal), as well as fine (ObF) and gross (ObG) actions, shown separately for the hemisphere contralateral (left column) and ipsilateral (right column) to the hand used. Solid lines depict group means for the comparison (green) and DCD (purple) groups; shaded regions indicate ±SEM across participants. Vertical dashed lines mark task onset (0 s) and offset (10 s); the horizontal dashed line indicates baseline (0%). Colored asterisks denote significant within-group deviations from baseline (one-sample permutation tests, $p < 0.05$).
